# Neptune: An environment for the delivery of genomic medicine

**DOI:** 10.1101/2021.01.29.428608

**Authors:** Eric Venner, Victoria Yi, David Murdock, Sara E. Kalla, Tsung-Jung Wu, Aniko Sabo, Shoudong Li, Qingchang Meng, Xia Tian, Mullai Murugan, Michelle Cohen, Christie Kovar, Wei-Qi Wei, Wendy K. Chung, Chunhua Weng, Georgia L. Wiesner, Gail P. Jarvik, Donna Muzny, Richard A. Gibbs, eMERGE Consortium

## Abstract

**Purpose:** Genomic medicine holds great promise for improving healthcare, but integrating searchable and actionable genetic data into electronic health records remains a challenge. Here, we describe Neptune, a system for managing the interaction between a clinical laboratory and an electronic health record system.

**Methods:** We developed Neptune and applied it to two clinical sequencing projects that required report customization, variant reanalysis and EHR integration.

**Results:** Neptune enabled the analysis of data for generation of and delivery to EHR systems of over 15,000 clinical genomic reports. These projects demanded customizable clinical reports that contained a variety of genetic data types including SNVs, CNVs, pharmacogenomics and polygenic risk scores. Two variant reanalysis activities were also supported, highlighting this important workflow.

**Conclusions:** Methods are needed for delivering structured genetic data to EHRs. This need extends beyond developing data formats to providing infrastructure that manages the reporting process itself. Neptune was successfully applied on two high-throughput clinical sequencing projects to build and deliver clinical reports to EHR systems. The software is open and available at https://gitlab.com/bcm-hgsc/neptune.

## Introduction

The goal of genomic medicine is to improve clinical outcomes^1^ by identifying patients at higher risk for specific adverse drug events, providing molecular diagnoses for etiologically heterogeneous diseases and identifying patients at increased lifetime risk of genetic disease. However, its implementation is currently limited by many factors. These include: 1) a lack of infrastructure for high-throughput clinical reporting^2–4^, 2) regulatory requirements for handling protected health information (PHI)^5,6^, 3) an expensive and manual process for genomic variant interpretation^7^, 4) data integration requirements that can rely on proprietary data formats that are different for each clinical site^8,9^ 5) a low frequency of actionable findings in some disease areas^10^, 6) an additional burden on providers to integrate genetic data in their practice^11^ and 7) a reluctance from insurance providers to cover the expense of tests underlying precision medicine^12^. All of these challenges demand research utilizing large genomic datasets connected with clinical outcomes measures to enable broad implementation of genomic medicine. Many national and international clinical sequencing projects have been established to fill this need, including the eMERGE Network^13^, All of Us (joinallofus.org), the IGNITE network^14^, and the Clinical Sequencing Evidence-Generating Research^15^ (CSER) consortium as well as a large number of private and regional initiatives^16,17^.

Integrating genomic data in electronic health records (EHRs) will help address some of these challenges. These data allow researchers to understand and improve the long-term clinical impact of genomic data and to demonstrate its utility to outside groups. Further, adding genetic data in a structured format to the EHR allows it to be searchable by clinicians. Unfortunately, genomic data are often heterogeneous, mix or lack standards, are updated regularly, and require domain expertise to handle correctly, with numerous edge cases. Although data standards are in development (https://emerge-fhir-spec.readthedocs.io/en/latest/), there is currently a lack of flexible, comprehensive, and open solutions for structuring genomic data and cleanly bridging the gap to EHR systems.

In support of the goal of delivering genomic data to the EHR, we have developed Neptune, an environment that enables users to identify and report known disease-causing variants in gene sets of interest, to gather curations for potential novel pathogenic variants from an external review system, and to share and update structured genomic knowledge with EHR systems and collaborators. An expert reviewer, upon receiving quality-controlled sample data, is presented with a highly filtered list of genomic variants to interpret according to ACMG guidelines^18^. Once variant review is complete, reports are automatically generated and made available for approval by a laboratory director. For applications which require handling of protected health information (PHI), our system is both capable of running in a HIPAA-compliant environment and operating as a hybrid system, with non-PHI elements running outside of the HIPAA-compliant environment.

Genomic variants are often interpreted as having ‘unknown significance’, especially for novel findings in non-European individuals. However, over time, new evidence emerges and these variants can be reclassified as either pathogenic or benign. Neptune supports the variant reanalysis workflow by tracking updated variant classifications in the VIP database, managing the generation of updated reports, and tracking report versions by integrating with the ARBoR^19^ system.

The key features of Neptune are: 1) to take as input genomic data (genotypes and coverage information) and compare against a ‘VIP database’ of known genetic variation, marking known variants with previously-curated data and selecting novel genomic variants for review, 2) to combine data from diverse sources including sample metadata from a LIMS and variant information from the VIP database and output data in a structured report file ready to be accepted by EHR systems, 3) to convert that structured data into a customizable human-readable report, 4) to enable corrected and updated reports, and 5) to enable the reanalysis and re-interpretation of data over time. In this report we describe the workflow of the Neptune environment, and describe its use for the implementation and customization of the clinical reporting workflow for two large genomic data integration into clinic projects: eMERGE III and HeartCare.

## Material and Methods

The primary functionality of Neptune begins after a list of genomic variants for each sample is created using standard bioinformatics pipelines^20^. The design of Neptune allows fully-automated reporting provided that all of the genomic variants present in a sample have been previously curated and stored in the ‘VIP’ variant database. Annotated variants and associated metadata are used to populate a structured .json format that represents the ‘clinical report’ for that sample. This functionality is encapsulated in an API (Table 1) to support interaction with external systems.

**Table 1.**

| Command | Input(s) | Description |
| --- | --- | --- |
| annotate | vcf, project configuration | Annotates vcf with data from the VIP. Marks if a variant has been previously seen or needs review |
| renderPreReport | VIP annotated vcf, configuration information for external data, report template, project configuration | Loads data from external sources, creates a structured output files |
| renderFinalReport | Pre-report file, project configuration | Populates the pre-report with the final set of data. Separate in case it needs to run in a PHI environment |
| reanalyze | VIP annotated vcf | Takes an existing VIP annotated vcf, shows differences to current VIP |

### VIP Database

Central to the Neptune process is the ‘VIP Database’ of genomic variation. The VIP Database contains variant information (chromosome, position, reference allele, variant allele), minor allele frequencies, transcript data (genomic variant effect, amino acid change), gene annotations (disease association, inheritance) and a large set of internal curation data (pubmed ids of related publications, comments and categories from clinical sites). It currently contains 381,564 variants (Figure 1B). This database was initially seeded by the two clinical reporting laboratories for the eMERGE III network^21^, and has been subsequently updated for novel variants that are detected in samples in the HGSC Clinical Lab and other public variant resources. This resource draws on both public resources (ClinVar, OMIM, literature review) and internal data sets. The VIP database is available for download at https://gitlab.com/bcm-hgsc/neptune.

**Figure 1:**
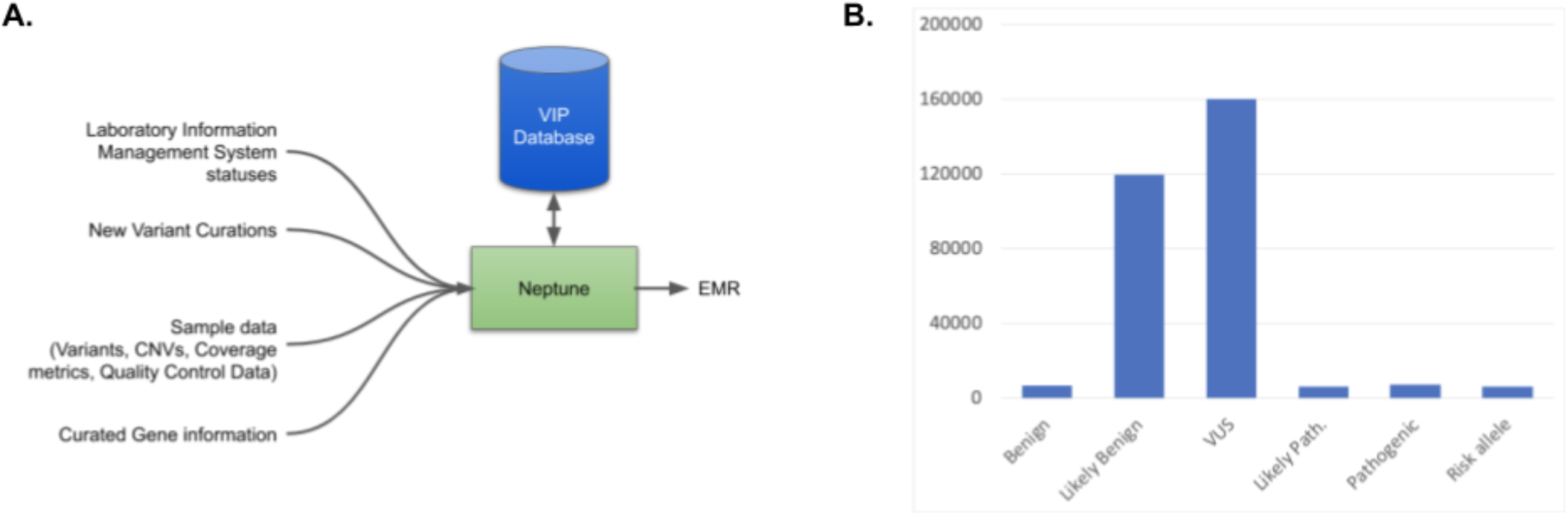
Overview of Neptune functionality. A. Neptune manages the variant review process and brings together disparate data from multiple external systems in order to create a final report file, in either json, html or FHIR format. Central to this process is the ‘VIP’ database of genetic variation. For each sample, novel genomic variants are added to this database and curated as needed according to project-specific rules. B. The contents of the VIP database includes curated variants. VIP database variants are predominantly VUS or Likely Benign.

### Variant Filtering and Annotation with Locally-Curated Variant Data

Neptune interacts with a snapshot of the VIP database in vcf format. Novel variants in a sample are detected by comparing their genomic coordinates and alternate allele to those in the database. Variants that are not present in the VIP database can be forwarded to a variant review system for manual curation, but in general, manual review of variants is the exception. In the eMERGE and HeartCare projects, over 99.99% (682343/682398) from a representative sample) of variants were handled automatically. Following manual curation, novel variants are added to the VIP database. Once all variants in a sample have been categorized, Neptune extracts reportable, pathogenic variants using curations stored in the VIP database, and outputs an automated clinical report populated with prioritized variants (or a negative report if no relevant variants are found).

The assessment of variants reviewed per sample in this study (Figure 2) was done by “replaying” our review process, starting from an empty VIP database. Variants were limited to the 68 eMERGE consensus reportable genes (*BMPR1A, CACNA1A, COL5A1, HNF1A, HNF1B, KCNE1, KCNJ2, OTC, PALB2, POLD1, POLE, SMAD4, ACTA2, ACTC1, APC, APOB, BRCA1, BRCA2, CACNA1S, COL3A1, DSC2, DSG2, DSP, FBN1, GLA, KCNH2, KCNQ1, LDLR, LMNA, MEN1, MLH1, MSH2, MSH6, MUTYH, MYBPC3, MYH11, MYH7, MYL2, MYL3, MYLK, NF2, PCSK9, PKP2, PMS2, PRKAG2, PTEN, RB1, RET, RYR1, RYR2, SCN5A, SDHAF2, SDHB, SDHC, SDHD, SMAD3, STK11, TGFBR1, TGFBR2, TMEM43, TNNI3, TNNT2, TP53, TPM1, TSC1, TSC2, VHL, WT1*). Each sample was analyzed in the order in which it was received. For each variant selected for review during our initial review process, we checked whether it was previously added to the database. For the first few samples analyzed in this ‘replay’ process, the database was empty or nearly empty, and therefore many variants needed to be assessed. Following the review, we added all reviewed variants, along with their variant classification, to the database, with the variant classification arrived at during the initial review. As we progressed through each of the 7258 samples from the data freeze, we recorded how many reviewable variants were not present in the database for each new sample.

**Figure 2:**
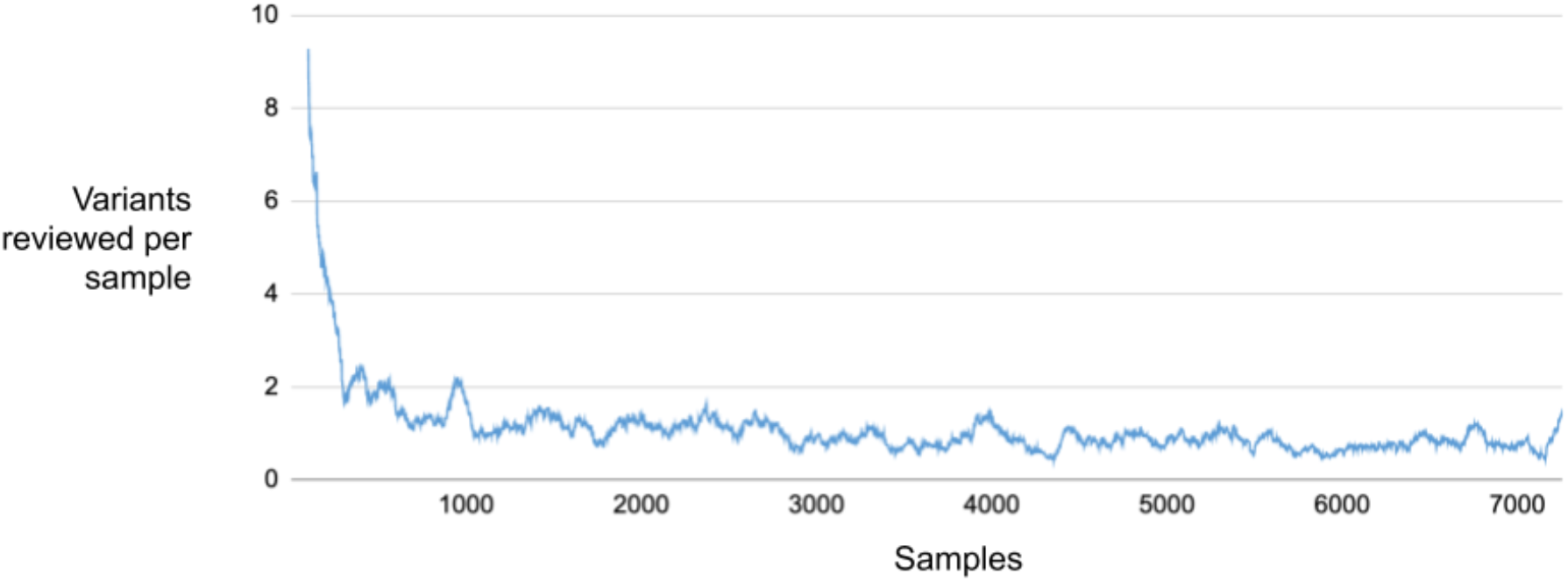
Variant Review Burden Over Time. The plot shows the number of variants per sample requiring review in 68 eMERGE III consensus reportable genes, starting with an empty database. As additional samples are reviewed from a data freeze of 7258, the number of variants per sample that are selected quickly decreases. In eMERGE III, the number of variants that require review plateaus at around 1 variant per sample.

### Copy-Number Variation

Neptune has an option of integrating copy number variants (CNVs) by incorporating AtlasCNV^22^ output into the reporting pipeline. AtlasCNV identifies copy number variants for each exon target of the capture design within a capture midpool experiment. The sample with median read depth at each exon target of interest is used as the reference sample, and a RPKM (sample normalization by Reads Per Kilobase per Million) log2 score (sample/reference ratio) is computed for each sample for a given exon target. Cutoffs are then applied to classify the CNV as either a heterozygous deletion, a homozygous deletion or duplication, and QC metrics flag regions where coverage is too variable to accurately call CNVs. If activated, reports contain a CNV section, alongside SNP and indel variants. CNVs and SNVs are reported alongside one another to highlight cases of compound heterozygosity, in which one gene contains both a CNV and another deleterious variant.

### Pharmacogenomics

Pharmacogenomics analysis is available for a subset of commonly reported genotypes and star alleles^23^. The module is configurable and the set of reported pharmacogenomic findings that are reported are defined using a mapping file that links reportable genotypes to their associated star alleles, phenotypes, and interpretation notes. Pharmacogenomic analysis requires either a gvcf input or external QC file with coverage values for all pharmacogenomic variant sites. Variants are assumed to be unphased, leading to ambiguous star allele assignments in some cases (e.g. TPMT *1/*3A vs *3b/*3c). If the pharmacogenomic analysis is active, an additional table will be added to the report that describes the pharmacogenomic variants in the patient, as well as adding the corresponding data to the structured JSON file.

### Polygenic Risk Scores

We developed a PRS module in Neptune to enable the clinical reporting of PRS, which will hopefully facilitate the gathering of clinical datasets that can be used to assess their utility. The polygenic risk score module reads a file in variant call format (vcf), restricted to sites of interest for a given polygenic risk score. It then calculates the risk score, using weights provided in a configuration file and the zygosity of each allele. This configuration file contains the set of sites to use for calculating the score, with the weight to use for each site if it is present in the sample. Lastly, the score for each sample is then compared against a reference distribution to determine the risk category for that sample.

### Report Templates

Reports are designed to meet all CAP / CLIA requirements and are highly customizable using an html-based templating system. Sections of the report can be activated or deactivated based on sample metadata such as project or sequencing methodology. Neptune supports both corrections and amendments to existing reports, with changes tracked and timestamped. By integrating with our variant review system, our internal deployment of Neptune streamlines the generation of batches of negative reports, which is critical in projects with a large number of negative reports.

### Conversion to Structured Data Formats

Neptune allows structured outputs to be in one of a variety of formats, including FHIR, HTML and JSON. Regardless of the format, the output captures all elements of the report including variant information, descriptive text, and coverage statistics produced by the ExCiD software. In the next step, this ‘pre-report’ is merged with PHI within a fully HIPAA-compliant environment and the final report is made available to a laboratory director for approval. For ease of viewing, an html version of the report is also made available.

For the eMERGE III project, the JSON file was converted into a proprietary XML format selected for use by the eMERGE network. This format was standardized across the two clinical reporting laboratories which allowed clinical sites to accept reports in a unified format^24^. In our HeartCare project, work is ongoing to develop a FHIR-compatible data specification and a conversion tool that can take this specification and JSON data to produce FHIR-compatible outputs (https://emerge-fhir-spec.readthedocs.io/en/latest/).

### Report Tracking and Verification with ARBOR

In order to track and verify report updates, Neptune integrates with the ARBoR system^19^. When the module is active, a unique barcode is added to the report. Our deployment of Neptune uses an instance of the ARBoR service, allowing it to write to an encrypted ledger of report entries each time a report is signed-out. The ARBoR client can then scan the barcode and verify the report against the ledger, alerting the user if there is a more recent report. The ledger does not require a database to function, so it persists beyond the lifespan of a particular project and does not require active maintenance.

### The BCM HeartCare study

In the Baylor College of Medicine (BCM) HeartCare study, patients who presented at BCM clinical sites were invited to participate in a clinical genomics study that included return of genomic results and integration into the EHR. This project increased the overall complexity of the clinical report by adding a section for reporting a polygenic risk score alongside integrated small variant and copy number variant genomic findings from 168 genes related to cardiac disease, pharmacogenomic findings for a set of drugs related to cardiovascular disease, and the reporting of two risk alleles for LPA^25^.

## Results

### Case Study: Electronic Medical Records and Genomics Network

The eMERGE Network brings together researchers and clinical laboratories from across the United States to investigate and improve systems for the implementation of genomic medicine^21^. Previously, as part of the eMERGE III Network, we performed clinical interpretation and issued over 14,500 clinical reports to 7 clinical sites for a targeted-gene panel of 68 consensus genes with additional clinical site specific genes. One particularly challenging aspect of this project was the requirement that clinical reports be highly customized for the needs of each clinical site. These customizations included modifying the gene list depending on the clinical site, allowing specific SNPs to be reported depending on the clinical site, adding a polygenic risk score section for one clinical site and hiding it from others, displaying a pharmacogenomic section for some sites and modifying the content of that section depending on site preferences, and modifying which set of metadata was displayed depending on the clinical site. Neptune implemented these customizations by employing a templating system that can key off sample-specific metadata that is pulled from the LIMS.

Genomic variants were interpreted according to ACMG guidelines^18^ in a high-throughput manner that relied on a set of automated filters, defined prior to the project start. Taking advantage of recurrent variant interpretations using the VIP database, we observed a rapid decline in novel variants per sample, followed by a stabilization around one reviewable variant per sample (Figure 2). A key lesson-learned was the benefit of gene-centric reviews; we adopted a review approach that ‘batched’ together a large number of samples (typically 1,200), and then reviewers would curate all variants in a particular gene from this batch in a single session. For example, a typical batch might contain 10 rare *BRCA2* variants; these would all be interpreted in the same session by one reviewer. This approach reduced context switching for reviewers and streamlined literature review. For example, large functional studies might encompass several variants from this review set.

As part of eMERGE III, we engaged in multiple reanalysis activities, supported by Neptune. Using an add-on system called ‘REVU’ (Reanalysis of Variants and Updater) we compared two snapshots of the ClinVar download (available from ftp://ftp.ncbi.nlm.nih.gov/pub/clinvar/), one from August 2018 and one from August 2019. Variants with a new Pathogenic or Likely Pathogenic interpretation where there was none previously were considered candidate ‘upgrades’. Any variants where a previous Pathogenic assertion had been removed, leaving only VUS, Benign or Likely Benign, was a candidate for a classification ‘downgrade’. For variants present in our eMERGE III cohort, we performed a re-interpretation. In the genomic regions covered by our test, we identified 614 unique variants with changed assertions. We next limited the search to reported genes for upgrades and reported variants for downgrades, resulting in 109 unique variants to review (99 upgrades, 10 downgrades) of which 34 (28 upgrades, 6 downgrades) of these were ClinVar ‘2 star’ variants and above. For each of these variants, we performed a full, manual variant interpretation, considering all ACMG evidence categories. Ultimately, we found five variants with sufficient evidence to change the variant interpretation and issued corrected reports. As an example, the NM_001127328.1:c.997A>G variant in *ACADM* (clinvar accession VCV000003586.19) was flagged for reanalysis due to one VUS interpretation that was submitted in April 2019. However, there are also 22 pathogenic or likely pathogenic interpretations that have been submitted to ClinVar for this variant, and in our assessment the variant remains pathogenic by applying ACMG PS3 (Well-established functional studies show a deleterious effect) and PS4 (Prevalence in affecteds statistically increased over controls) subcategories. In future efforts we will assess alternate filtering strategies that may help eliminate cases like this. The total time required for manual review varied greatly from between a few minutes and many (> 5) hours, based primarily on the additional information available about the variant and the number of discussions required by the review team to finalize their interpretation. For first review, reanalysis took 32 minutes on average (std. dev 9.4). The majority of variants could be reclassified by a first reviewer, but a small fraction (< 9%) required attention from a laboratory director.

In a separate reanalysis study, we identified genomic variants of unknown significance (VUS) that could reach P/LP status with one additional ACMG sub-category, and prioritized these to gather additional phenotypic and family history information from clinical sites (Figure 3B). There were 83 variants identified initially, of which we reclassified 4, either using ACMG subcategory PS4 (prevalence in affecteds significantly increased over controls) or PP4 (patient’s phenotype or family history highly specific for gene). An example was the c.551T>C p.Leu184Pro (NM_000551.3) variant in the *VHL* gene, which was borderline VUS based on the evidence we had (PP3 - computationally predicted to be deleterious, PM2 - absent from population databases). Two papers reported the variant associated with affected individuals, but this was not enough evidence to apply PS4. However, upon contacting the clinical site, we learned that the patient was diagnosed with Von Hippel-Lindau disease, which allowed us to apply the PP4 subcategory, moving this variant to likely pathogenic.

**Figure 3:**
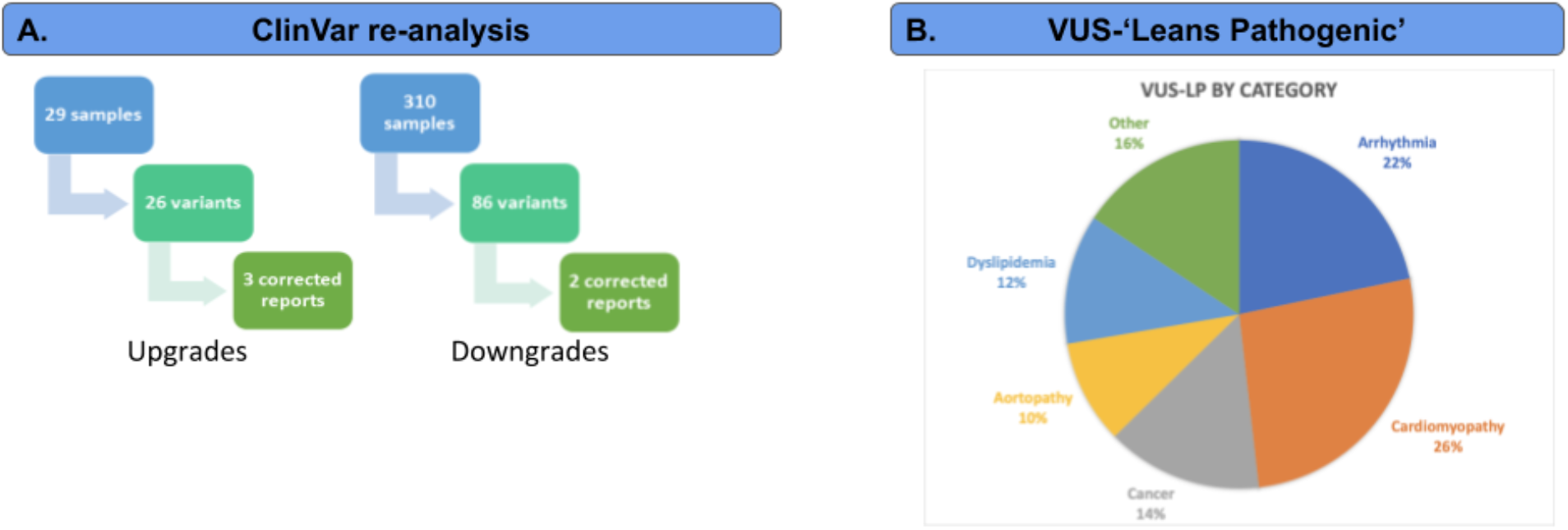
eMERGE III reanalysis activities. Neptune supported two parallel reanalysis activities during the eMERGE III project. First was a project with the goal of providing updated reports when variant classifications change (3A) over time. To accomplish this, we used Neptune’s reanalysis module to compare a ClinVar snapshot to local variant categorizations. We identified upgrades and downgrades by detecting either unreported variants with a new Pathogenic / Likely pathogenic classification in ClinVar or a reported variant with a new VUS, Benign or Likely benign classification. There were 26 upgrades for review, resulting in 3 updated reports (all initially VUS) and 86 downgrades for review, resulting in 2 updated reports. Next, we collected a set of VUS variants that were lacking one ACMG subcategory to reach an overall classification of likely pathogenic (3B). We then contacted clinical sites requesting more detailed patient phenotype information, in order to be able to apply the PP4 ACMG subcategory (Patient phenotype or family history highly specific for gene). In four cases we were able to issue updated reports, all due to the new clinical information. In a separate study, we reanalyzed 83 variants based on additional clinical information requested from clinical sites for variants that were VUS but which could be reclassified as Likely Pathogenic with the application of one ACMG subcategory. This resulted in four updated reports and highlights the importance of detailed clinical information during review by clinical geneticists.

In total, we re-issued nine reports based on variant classification updates. By using the size of the eMERGE panel (68 consensus genes) and the number of reports in circulation when we started that effort (approximately 15,000) we can estimate that the burden placed on clinical laboratories by reanalysis will require assessing 0.0001 (109 / 1,020,000) variants per gene on an issued report. The rate of reissued reports remains low, at 0.03% (5/15,000). As the number of interpreted variants increases, this problem will continue to grow.

### Case Study: BCM HeartCare

In a second application, we performed variant interpretation and reporting for 709 patients who presented at BCM cardiovascular clinics. 8.5% percent of the cases were positive for a pathogenic or likely pathogenic SNV or CNV, and 49% were positive for a pharmacogenomic finding. Management changes as a result of these findings included recommending additional specific laboratory testing including imaging, referral for a genetic consultation, or a change in medication.

The variant review requirements and workflows were similar to eMERGE, with one important difference: the addition of patient and family management recommendations, written by a clinical geneticist. This part of the report provided specific feedback to the ordering physician on managing a genetic finding, and when appropriate contained advice on additional testing, drug regimens to start or avoid, advice on additional genetic counseling, and recommendations on cascade testing. Composing the physician guidance section added significant amounts of time to report preparation. These changes were implemented by creating a new report template to support the additional fields. Figure 4 shows an example HeartCare report.

**Figure 4:**
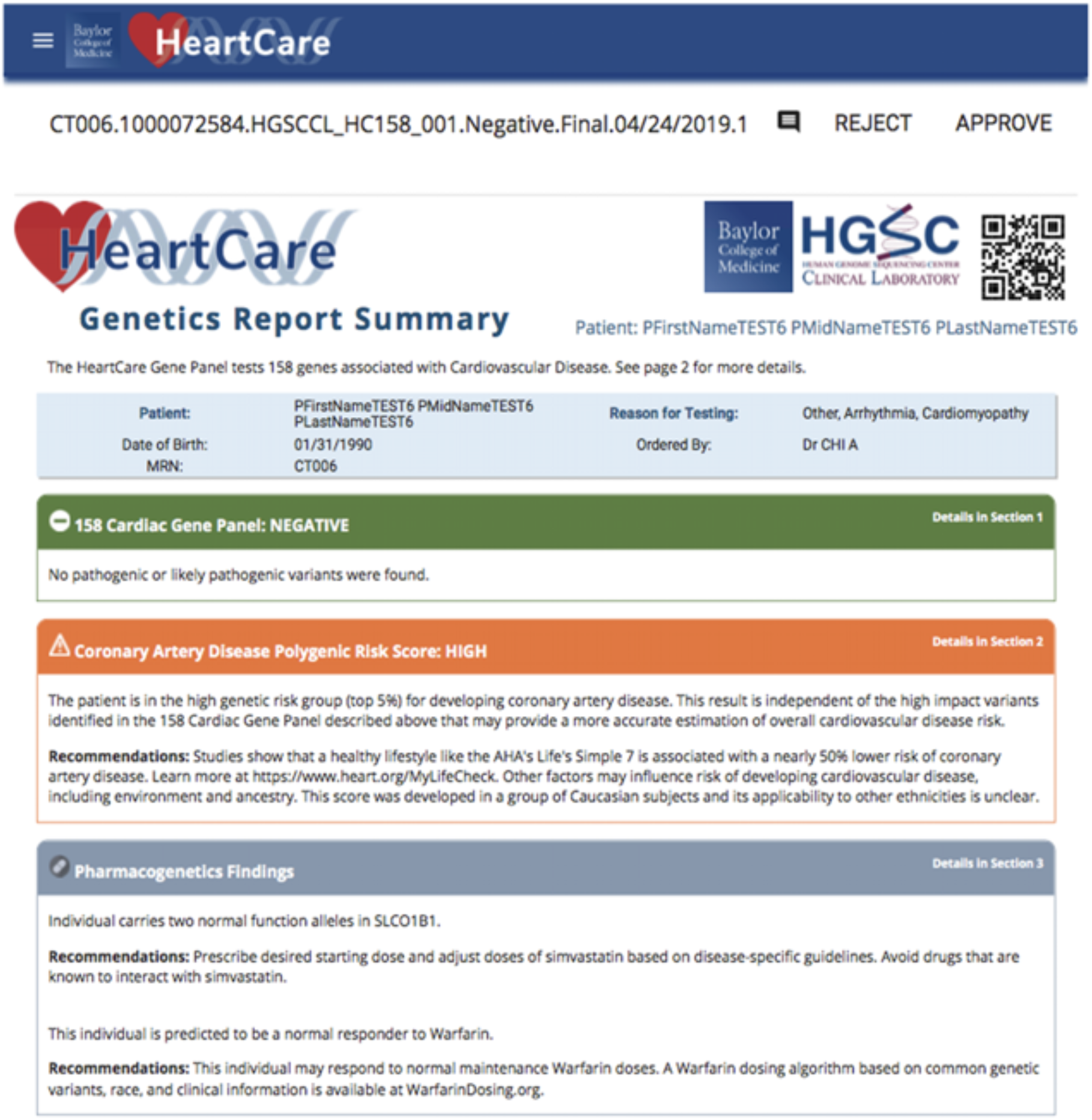
Example HeartCare report. The rendered report for the HeartCare cardiovascular disease report contains three sections: 1) findings from the set of 158 genes associated with cardiovascular disease, 2) Polygenic Risk Score findings and 3) Pharmacogenomic findings. Elements of the report are available in structure formats (json, HL7) and are fully rendered as a pdf. Reports were transmitted to patients and clinicians participating in our study via the EHR.

Another addition in HeartCare was the reporting of polygenic risk scores. We employed a previously developed polygenic risk score for coronary artery disease (Khera et al., 2016) which previously demonstrated a difference in hospitalization between high and low risk groups, finding that high risk individuals have a 91% higher relative risk than low risk individuals. For our assay, after clinician feedback, we chose a stringent threshold for high risk and reported the top 5% of individuals in this distribution as the “high risk” group (Top 5% >= 4.5824).

We also implemented a HIPAA-compliant reporting portal, hosted on AWS, for the final report rendering and storage. In addition, we piloted an EPIC integration feature and built an HL7v2 message generator that will be integrated into Neptune in a future version. This allowed HeartCare reports to be integrated into the EHR in a structured format, instead of simply loading a report in pdf format. A full description and lessons learned from the HeartCare study are described in Murdock et al. 2020 (in preparation).

## Discussion

Neptune provides a highly customizable platform that enables the delivery of genomic results to support genomic medicine. It facilitates complex reporting workflows including reanalysis, and connects genomic data to clinical geneticists and the EHR, enabling flexible, customized reporting. It is backed by a VIP database of genetic variation that stores the updated variant curations. We have deployed this environment to enable two exemplar projects in which clinical genetic data were reviewed, reported out and transferred back to a clinical site. Neptune is a validated approach to clinical genetic reporting that can alleviate some of the problems related to delivering scalable clinical genetic data.

Reanalysis places a substantial workload on clinical genetics activities and the overall effort will increase with the volume of reports issued. Based on the number of genes present on the gene panel designs used in the tests reviewed here, we observed a rate of 0.0001 variants per gene on an issued report per year. Thus, when reporting clinical genetic data at a large scale, complete reanalysis may not be feasible and clear guidelines will be crucial to define the extent to which reanalysis activities are necessary. Future work will examine the extent to which accelerating submissions to ClinVar might change this estimate and whether potential increasing concordance between laboratories will reduce the amount of work remaining.

We and others have observed^26^ that a plateau is reached in the review burden per sample (Figure 2) as additional samples are added to the study. This highlights the ongoing and longterm difficulty of variant interpretation. Active efforts towards rule-based interpretation underway by ClinGen will help to address this problem; as more automatable genomic variant interpretation becomes standard, successively larger the projects will become feasible.

In many ways, the successful implementation of genomic medicine relies on structured integration of genomic data into the EHR systems, especially to enable clinical decision support (CDS). When stored in a structured format, these data can be acted on by CDS tools to provide context-dependent decision support to clinicians. Optimally, data would flow smoothly both into and out of the EHR. Health information can be used to support variant interpretation and genomic data are already proving actionable in the clinic, with its utility increasing rapidly. Data interchange formats like FHIR (https://emerge-fhir-spec.readthedocs.io/en/latest/) are crucial for enabling this interchange and will empower the next generation of clinical genomic integration.

## Data availability

The software is available from https://gitlab.com/bcm-hgsc/neptune

## Acknowledgements

Authors: Eric Venner^1,2^, Victoria Yi^1^, David Murdock^1,2^, Sara E. Kalla^1^, Tsung-Jung Wu^1^, Aniko Sabo^1,2^, Shoudong Li^1^, Qingchang Meng^1^, Xia Tian^1^, Mullai Murugan^1^, Michelle Cohen^1^, Christie Kovar^1^, Wei-Qi Wei^3^, Wendy K. Chung^4^, Chunhua Weng^5^, Georgia L. Wiesner^6^, Gail P. Jarvik^7^, Donna Muzny^1,2^, Richard A. Gibbs^1,2^,

## Author Information

Conceptualization: E.V., R.G.

Data curation: A.S., S.L., Q.M., X.T., M.C., D.M.

Formal Analysis:

Funding acquisition: R.G, D.M.

Investigation: E.V., M.C., D.M.

Methodology: E.V., R.G., V.Y., C.K.

Project administration: M.C., C.K., M.M.

Resources: W.W., W.C., C.W., G.W., G.J., R.G.

Software: E.V., V.Y., S.K., T.W.,

Supervision: E.V., M.M., D.M., C.K.

Validation: E.V., V.Y., T.W.

Visualization: E.V., M.C.

Writing – original draft: E.V., V.Y.

Writing – review & editing: E.V., V.Y., D.M., S.K., T.W., A.S., S.L., Q.M., X.T., M.M., M.C., C.K., W.W. W.C., C.W., G.W., G.J., D.M., R.G.

## Ethics Declaration

This work was funded by internal operating funds of the Baylor College of Medicine Human Genome Sequencing Center (HGSC), and by the NIH eMERGE program Phase III: U01HG8657 (Kaiser Permanente Washington/University of Washington); U01HG8685 (Brigham and Women’s Hospital); U01HG8672 (Vanderbilt University Medical Center); U01HG8666 (Cincinnati Children’s Hospital Medical Center); U01HG6379 (Mayo Clinic); U01HG8679 (Geisinger Clinic); U01HG8680 (Columbia University Health Sciences); U01HG8684 (Children’s Hospital of Philadelphia); U01HG8673 (Northwestern University); U01HG8701 (Vanderbilt University Medical Center serving as the Coordinating Center); U01HG8676 (Partners Healthcare/Broad Institute); and U01HG8664 (Baylor College of Medicine). The HGSC is a one of the two Sequencing Centers for the eMERGE III. The Electronic Medical Records and Genomics (eMERGE) Network is a National Human Genome Research Institute (NHGRI)-funded consortium tasked with developing methods and best practices for utilization of the electronic medical record (EMR) as a tool for genomic research. All 11 sample collection sites consented participants under Institutional Review Board (IRB)-approved protocols and the two sequencing centers had IRB-approved protocols that deferred consent to the participating sites. The protocol number for Baylor College of Medicine was (#H-40455).

## Notes

https://gitlab.com/bcm-hgsc/neptune

### Competing Interest Statement

Disclosure: E.V. is a cofounder of Codified Genomics, which provides variant interpretation
services. R.G., D.M., D.M., disclose that the Baylor Genetics Laboratory is co-owned by Baylor
College of Medicine. All other authors declare no conflicts of interest.

